# S phase R-loop formation is restricted by PrimPol-mediated repriming

**DOI:** 10.1101/318220

**Authors:** Saša Šviković, Alastair Crisp, Sue Mei Tan-Wong, Thomas A. Guilliam, Aidan J. Doherty, Nicholas J. Proudfoot, Guillaume Guilbaud, Julian E. Sale

## Abstract

During DNA replication, conflicts with ongoing transcription are frequent and require careful management to avoid genetic instability. R-loops, three stranded nucleic acid structures comprising a DNA:RNA hybrid and displaced single stranded DNA, are important drivers of damage arising from such conflicts. How R-loops stall replication and the mechanisms that restrain their formation during S phase are incompletely understood. Here we show *in vivo* how R-loop formation drives a short purine-rich repeat, (GAA)_10_, to become a replication impediment that requires the repriming activity of the primase-polymerase PrimPol for its processive replication. Further, we show that loss of PrimPol results in a significant increase in R-loop formation around the repeat during S phase. We extend this observation by showing that PrimPol suppresses R-loop formation in genes harbouring secondary structure-forming sequences, exemplified by G quadruplex and H-DNA motifs, across the genome in both avian and human cells. Thus, R-loops promote the creation of replication blocks at susceptible sequences, while PrimPol-dependent repriming limits the extent of unscheduled R-loop formation at these sequences, mitigating their impact on replication.

## Introduction

Tracts of repetitive sequence, known as microsatellites or short tandem repeats, occur frequently in vertebrate genomes (Clark et al., 2006, Tripathi & Brahmachari, 1991, Willems et al., 2014). Many such sequences are capable of forming secondary structures, including hairpins, cruciforms, triplexes (H-DNA) and G-quadruplexes (G4s), that have the potential to impede DNA replication (Mirkin & Mirkin, 2007). However, the factors that determine whether these sequences pose a barrier to DNA synthesis *in vivo* and the consequences of their doing so are not well understood.

It is well established that long repeats lead to problems with both replication and transcription. For example, a long tract of polypurine-polypyrimidine (GAA)_n_ repeats (in which n can exceed 1500) is linked to the inherited neurodegenerative disorder Friedreich’s ataxia (Campuzano et al., 1996). These repeats can form H-DNA (Frank-Kamenetskii & Mirkin, 1995) and are also prone to form R-loops (Groh et al., 2014), three-stranded nucleic acid structures in which nascent RNA hybridises to its complementary DNA template, displacing the non-template DNA strand (Thomas et al., 1976). These secondary structures block replication both in bacterial and yeast models, as well as in human cells (Chandok et al., 2012, Krasilnikova & Mirkin, 2004, Ohshima et al., 1998), which promotes genetic instability of the repeat (Gerhardt et al., 2016). They also perturb transcription by reducing RNA polymerase II (RNAPII) elongation (Bidichandani et al., 1998, Punga & Buhler, 2010) and lead to deposition of repressive chromatin marks (Al-Mahdawi et al., 2008, Saveliev et al., 2003). In the case of Friedreich’s ataxia this results in transcriptional silencing of Frataxin (*FXN*), in which the repeat is located.

Less clear is the impact of the much more common short repetitive tracts found throughout the genome (Clark et al., 2006, Willems et al., 2014). These have generally not been linked to any significant impact on replication or transcription. For example, the (GAA)_n_ repeat in normal alleles of *FXN* (n < 12) is not at risk of expansion (Schulz et al., 2009), despite the ability of even (GAA)_9_ to form a stable H-DNA structure at physiological pH *in vitro* (Potaman et al., 2004). Further, these ‘normal’ repeats also induce significantly less R-loop formation than disease-length alleles and are not associated with delay of RNAPII or transcriptional silencing (Groh et al., 2014). However, it remains unclear whether this apparently inert behaviour is due to these sequences being incapable of forming secondary structures or whether it is the result of activities that counter structure formation and its consequences.

In this paper, we address these questions by studying the replication of a short GAA repeat in the *BU-1* locus of chicken DT40 cells. We have previously used this approach to show that G-quadruplexes are able to impede the leading strand polymerase (Sarkies et al., 2012, Sarkies et al., 2010, Schiavone et al., 2014), and that repriming, performed by primase-polymerase PrimPol, is deployed frequently to mitigate such events (Schiavone et al., 2016). This is consistent with such structures forming a replication impediment even under unperturbed conditions (Schiavone et al., 2016). We now show that (GAA)_10_ requires PrimPol for its processive replication, demonstrating that these ubiquitous repeats pose an impediment to DNA synthesis. However, the ability of (GAA)_10_ to impede replication is entirely dependent on DNA:RNA hybrid formation as overexpression of RNaseH1 completely bypasses the requirement for PrimPol. Furthermore, loss of PrimPol promotes unscheduled R-loop accumulation around the (GAA)_10_ sequence during S phase and results in higher levels of R-loop formation in genes harbouring secondary structure-forming H-DNA and G4-motifs throughout the genome. These results provide a direct demonstration that R-loop formation can promote DNA sequences with structure forming potential to become replication impediments and that repriming by PrimPol, in turn, limits excessive R-loop accumulation in the vicinity of these sequences, we suggest by preventing excessive exposure of single stranded DNA during their replication.

## Results

### Instability of BU-1 expression monitors replication delay at (GAA)n

We have previously shown that expression instability of the *BU-1* locus in chicken DT40 cells provides a sensitive readout for replication delay at G4 motifs(Schiavone et al., 2014). The wild type locus contains a G4 motif 3.5 kb downstream of the TSS (the +3.5 G4) towards the end of the second intron (Fig 1A), which is responsible for stochastic, replication-dependent *BU-1* expression instability under conditions in which G4 replication is impaired (Guilbaud et al., 2017, Sarkies et al., 2012, Schiavone et al., 2014). Failure to maintain processive replication through the +3.5 G4 motif leads to uncoupling of DNA unwinding and DNA synthesis, interrupting normal histone recycling at the fork and the accurate propagation of epigenetic information carried by post-translational modifications on histone proteins (Fig 1A). This leads to replication-dependent instability of *BU-1* expression manifested as stochastic, replication-dependent conversion of the normal ‘high’ expression state to a lower expression level (Sarkies et al., 2012, Schiavone et al., 2014). This expression instability can be readily monitored by flow cytometric analysis of surface Bu-1 protein(Sarkies et al., 2012) (Fig EV1), providing a simple method to cumulatively ‘record’ episodes of interrupted DNA synthesis at the secondary structure-forming motif.

**Figure 1.**
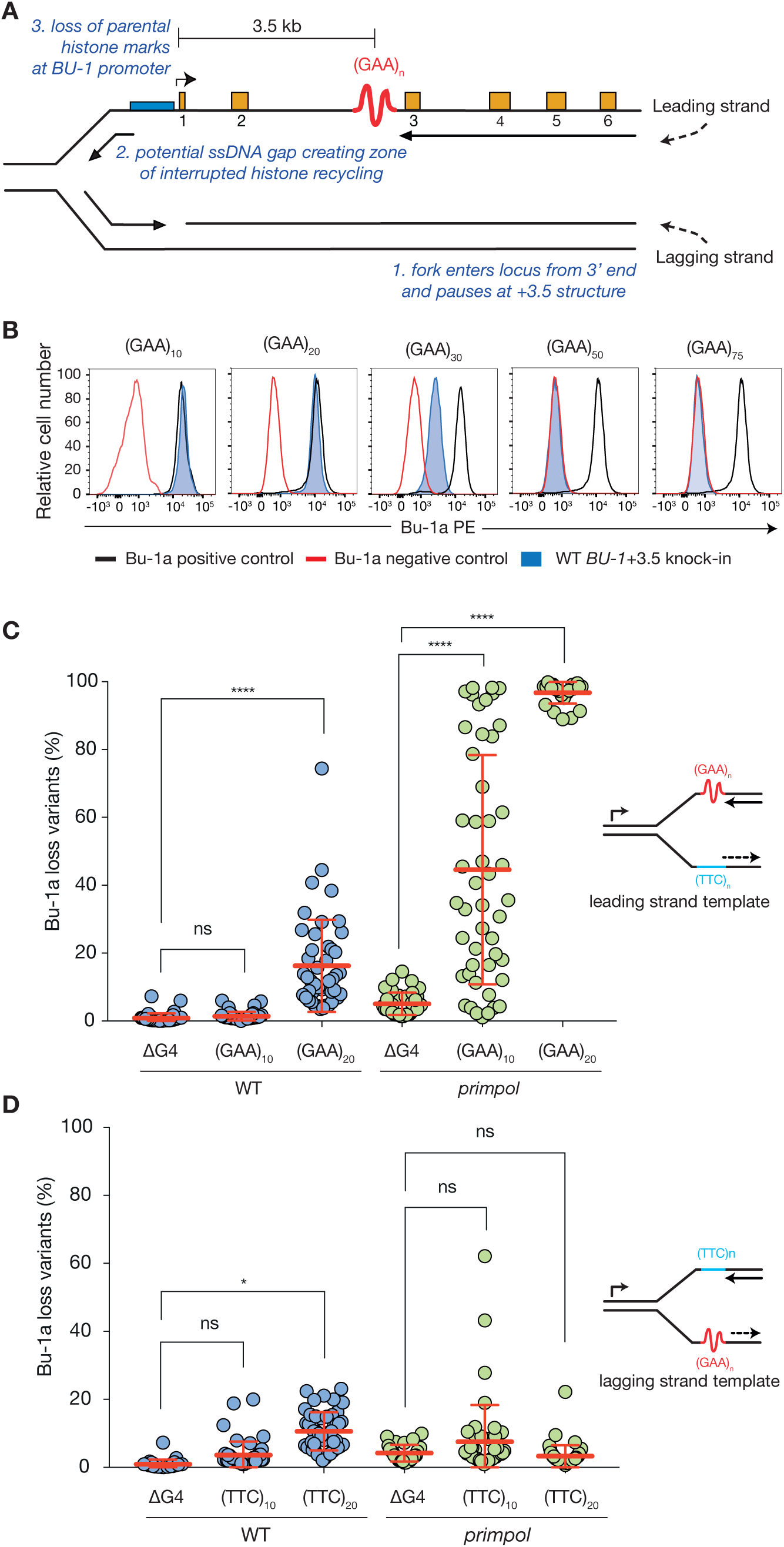
Short (GAA) tracts cause *BU-1* epigenetic instability in *primpol* cells. A. Expression instability of the chicken *BU-1* locus as a reporter for replication impediments formed by structure-forming DNA sequences. The leading strand of a replication fork entering the locus encounter a DNA sequence with structure-forming potential located 3.5 kb downstream of the transcription start site. In wild type cells this is a G4 motif, which is replaced by (GAA)_n_ repeats in this study. A polymerase stall leads to the formation of a putative ssDNA gap and loss of parental histone modifications in the zone of interrupted histone recycling. If this involves the promoter region it results in a change in expression of the gene. B. Flow cytometry for Bu-1a expression in wild type cells with (GAA)_n_ tracts of different length knocked into the *BU-1A* locus. DT40 cells are heterozygous and carry one *BU-1A* and one *BU-1B* allele. All experiments introducing repeats into *BU-1A* are carried out in cells in which the +3.5 G4 has been deleted from both *A* and *B* alleles, to avoid transvection between the alleles (Schiavone et al., 2014). Black outline: positive control; red outline negative (*BU-1* knockout) control. C, D. Bu-1a fluctuation analysis of wildtype and *primpol* cells in which the endogenous +3.5 G4 has been deleted (ΔG4) or with (GAA)_10_ and (GAA)_20_ sequence orientated such that it is replicated as the leading (C) or lagging (D) strand template for a fork entering from the 3’ end of the locus as shown in A. Circles represent percentage of Bu-1a loss variants in individual clones analysed, with mean ± SD reported. ****p < 0.0001, * p ≤ 0.05, ns = not significant; one-way ANOVA (in GraphPad Prism v.7.0).

To model the replication of (GAA)_n_ repeats we started with DT40 cells in which the +3.5 G4 motif of *BU-1* had been deleted on both the *A* and *B* alleles (Schiavone et al., 2014). (GAA)_n_ repeats of lengths between n = 10 and n = 75 were constructed either by synthesis for n ≤ 30 or, for the longer tracts, using a cloning strategy for highly repetitive sequences (Fig EV2). The repeats were then introduced into the *BU-1A* allele by gene targeting, as previously described (Schiavone et al., 2014) to create *BU-1A^(GAA)n^* cells. Following selection cassette removal, cells carrying (GAA)_10_ and (GAA)_20_ in *BU-1A* exhibited wild type expression levels (Fig 1b). However, the presence of (GAA)_30_ resulted in reduced expression of *BU-1A*, while in cells carrying (GAA)_50_ and (GAA)_75_, *BU-1A* expression was essentially abrogated (Fig 1b).

The reduced expression in cells carrying (GAA)_30-75_ affects the entire population and thus appears to be distinct from the stochastic, replication dependent loss of expression we have previously reported to be induced by G4 motifs in cells lacking enzymes involved in G4 replication (Sarkies et al., 2012, Sarkies et al., 2010, Schiavone et al., 2014). Rather, these longer repetitive tracts resulted in the accumulation of chromatin-associated nascent RNA (ChrRNA) (Nojima et al., 2016) within the locus (Fig EV3a), consistent with impaired expression being due to reduced processivity of RNA polymerase II. As the global reduction of *BU-1* expression in (GAA)_30-75_ alleles precluded the analysis of stochastically generated loss variants, we focussed our subsequent analyses on (GAA)_10_ and (GAA)_20_.

Fluctuation analysis for the generation of Bu-1a loss variants confirmed that the presence of (GAA)_10_ at the +3.5 kb position did not affect the stability of *BU-1* expression in a wild type background (Fig 1c). However, (GAA)_20_ induced modest, but significant, formation of Bu-1a loss variants (Fig 1c) suggesting that this repeat is able to impede replication even in wild type conditions. We next examined the effect of deleting PrimPol to examine the extent to which repriming mitigates the replication impediment posed by these sequences. The results were striking: the rate at which Bu-1a loss variants were generated in *primpol* cells increased significantly, both for (GAA)_10_ and (GAA)_20_ (Fig 1c). These observations are consistent with repriming preventing significant uncoupling of DNA unwinding and DNA synthesis at these short repeats. They also suggest that these sequences are forming frequent impediments to otherwise unperturbed DNA replication.

The *BU-1* instability seen in *primpol* cells carrying (GAA)_10_ is comparable to that observed in *primpol* cells harbouring the endogenous +3.5 G4 (Schiavone et al., 2016). Similar Bu-1a^medium^ and Bu-1a^low^ expression states, characterised by loss of H3K4me3 and additional DNA methylation respectively, were isolated (Fig EV3b-e). Genetic instability, at a level that could explain the formation of Bu-1a loss variants, was not detected (Fig EV3f). Together, these observations suggest that the (GAA)_10_ sequence causes epigenetic instability through the same replication-dependent mechanism that we have previously described for G4s. The effect of the repeat was orientation dependent (Fig 1d), only producing instability when knocked-in such that the purine-rich strand formed the leading strand template for a fork entering from the 3’ end of the locus.

### The RPA-binding and repriming functions of PrimPol are required to ensure processive replication at the BU-1 (GAA)_10_ repeat

While PrimPol can perform some translesion synthesis, considerable evidence now supports repriming as its main *in vivo* role (Keen et al., 2014, Kobayashi et al., 2016, Mouron et al., 2013, Schiavone et al., 2016). The repriming function of PrimPol requires the C-terminal zinc finger and RPA binding motif A (RBM-A), which mediates an interaction with the single-stranded binding protein replication protein A (Guilliam et al., 2017, Guilliam et al., 2015, Wan et al., 2013). To confirm that the primary *in vivo* role played by PrimPol in the replication of (GAA)_10_ is indeed repriming, we performed a complementation study by ectopically expressing YFP-tagged human PrimPol in *primpol* cells (Fig 2). Expression of full-length human PrimPol completely restored the stability of *BU-1A* expression in *primpol* cells carrying the (GAA)_10_ repeat. Expression of catalytically inactive PrimPol (D114A, E116A or hPrimPol[AxA]) and the DNA-binding zinc finger mutant (C419A, H426A or hPrimPol[ZfKO]) shows that neither is able to prevent instability of *BU-1* expression (Fig 2), a result concordant with our previous observation in cells harbouring the natural +3.5 G4. Importantly, RBM-A, but not RBM-B, of PrimPol is essential for efficient suppression of (GAA)_10_ repeat-induced instability of *BU-1* expression, consistent with previous work showing that while both RBM-A and RBM-B can bind the same basic cleft in RPA70N *in vitro*, only RBM-A appears important *in vivo* (Guilliam et al., 2017).

**Figure 2.**
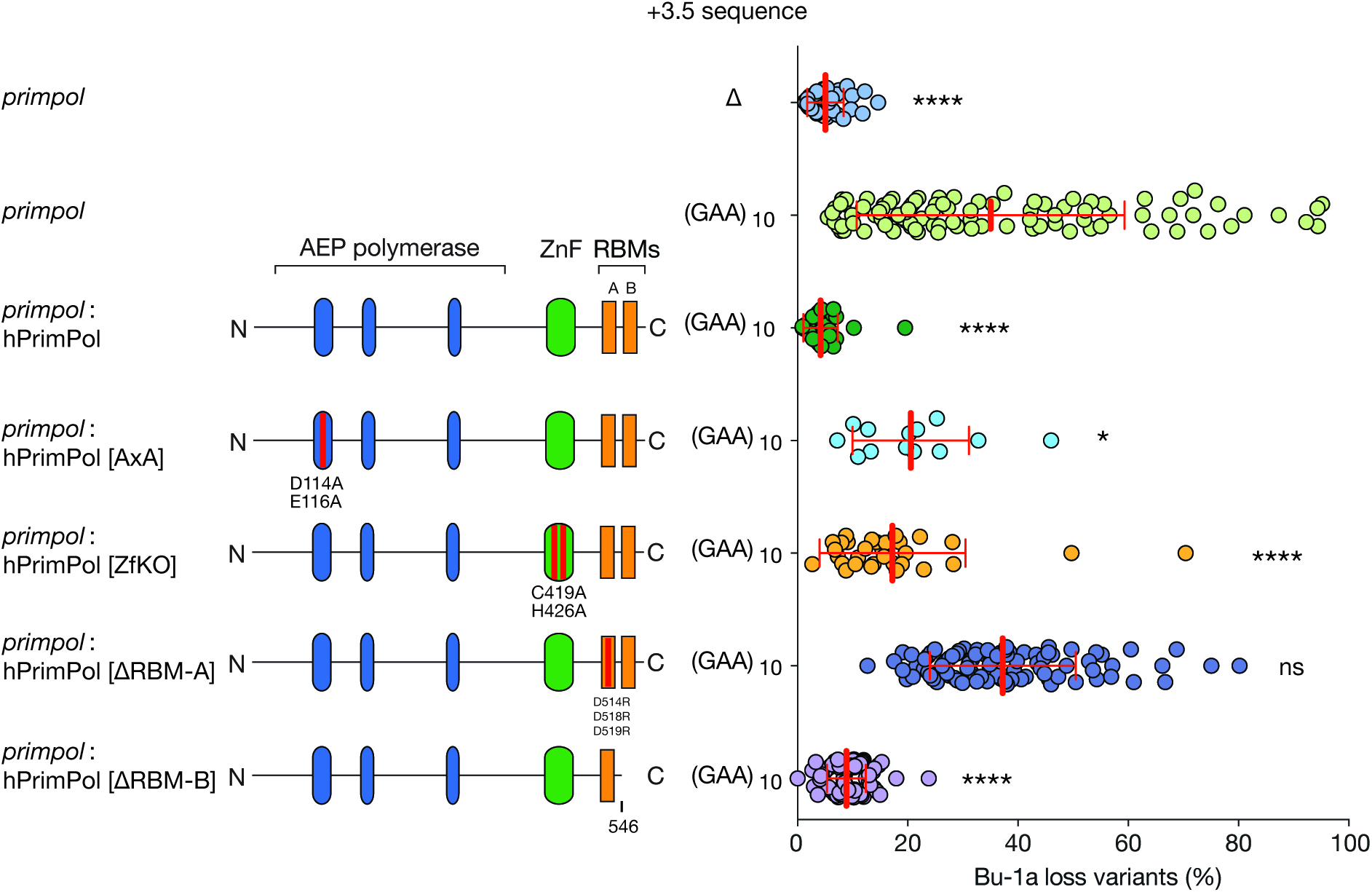
The repriming function of PrimPol is required to maintain expression stability of *BU-1* harbouring a (GAA)_10_ repeat. Human PrimPol, or mutants, tagged with YFP were expressed in *primpol* cells harbouring (GAA)_10_ sequence in the *BU-1A* locus. Bu-1a and YFP double positive cells were sorted and expanded for 2 weeks, and then analysed for Bu-1a expression variants. For each complementation, at least two independently derived clones were subjected to fluctuation analysis. As previously observed (Schiavone et al., 2016), expression of hPrimPol[AxA] and hPrimPol[ZfKO] is deleterious and unstable. Cells expressing these mutations and remaining YFP-positive at the end of the expansion period will have been through fewer divisions the other lines in this analysis. Pooled results (mean ±SD) are represented here. Statistical significance: **** p < 0.0001, * p ≤ 0.05, ns = not significant; one-way ANOVA.

REV1, a Y-family DNA polymerase, is required for maintaining stability of *BU-1* expression when the sequence at the +3.5 kb position is a G4 motif (Sarkies et al., 2012, Schiavone et al., 2014), which reflects a direct role for REV1 in G4 replication (Eddy et al., 2014, Sarkies et al., 2010). Replacing the +3.5 G4 in *rev1* cells with (GAA)_10_ repeats did not result in significant destabilisation of *BU-1* expression (Fig EV4a), while (GAA)_20_ results in a modest destabilisation of *BU-1* expression, comparable to the effect in wild type cells. This suggests that the role REV1 plays in maintaining epigenetic stability of *BU-1* is specific to G4 motifs in contrast to PrimPol, which is involved in replicating both types of secondary structure through its ability to reprime.

### PrimPol limits R-loop formation around a (GAA)_10_ repeat

The orientation dependence of the GAA tract with respect to *BU-1* instability is in line with the predicted formation of triplex DNA when a polypurine tract is transcribed as the coding strand, but not as the template strand (Grabczyk & Fishman, 1995, Grabczyk et al., 2007). *In vitro* studies have shown that the formation of triplexes at (GAA)_n_ occurs concurrently with the formation of a stable DNA:RNA hybrid between the TTC-rich template strand and the nascent GAA-containing RNA strand (Grabczyk et al., 2007). Furthermore, pathological formation of R-loops has been reported at long (GAA)_n_ repeats (n ≥ 650) in immortalised lymphoblasts derived from Friedreich’s ataxia patients (Groh et al., 2014). These reports, together with the results presented thus far, prompted us to investigate whether R-loops contribute to replication stalling induced by short GAA tracts in *BU-1*.

R-loops can be detected using a DNA:RNA hybrid-specific antibody, S9.6 (Boguslawski et al., 1986). We first examined R-loop formation in the (GAA)_10_-containing *BU-1* locus of wild type cells using DNA:RNA immunoprecipitation (DRIP) followed by quantitative PCR (Fig 3a). The first striking feature to note is the strong signal in the vicinity of the transcription termination site (Fig 3a), consistent with the previously described R-loops formed in this region in a subset of genes that are mechanistically associated with transcription termination (Skourti-Stathaki et al., 2014). Interestingly, this signal is consistently higher in *primpol* cells compared with wild type, although we do not have a clear explanation for this observation at present. In the body of *BU-1* in wild type, the presence of (GAA)_10_ at the + 3.5kb position results in a very modest DRIP signal in the vicinity of the repeat. In contrast, *primpol* cells exhibit a highly significant increase in R-loop signal around the same repeat. Crucially, this enrichment is abrogated when the repeat is deleted, showing that R-loop accumulation is not simply a direct consequence of loss of PrimPol, but arises due to failure of repriming at the replication impediment created by (GAA)_10_. This is unlikely to reflect a direct role of PrimPol in processing R-loops as *primpol* cells are not sensitive to camptothecin, a topoisomerase I inhibitor that leads to increased R-loop formation (Fig EV5) (Kobayashi et al., 2016).

**Figure 3.**
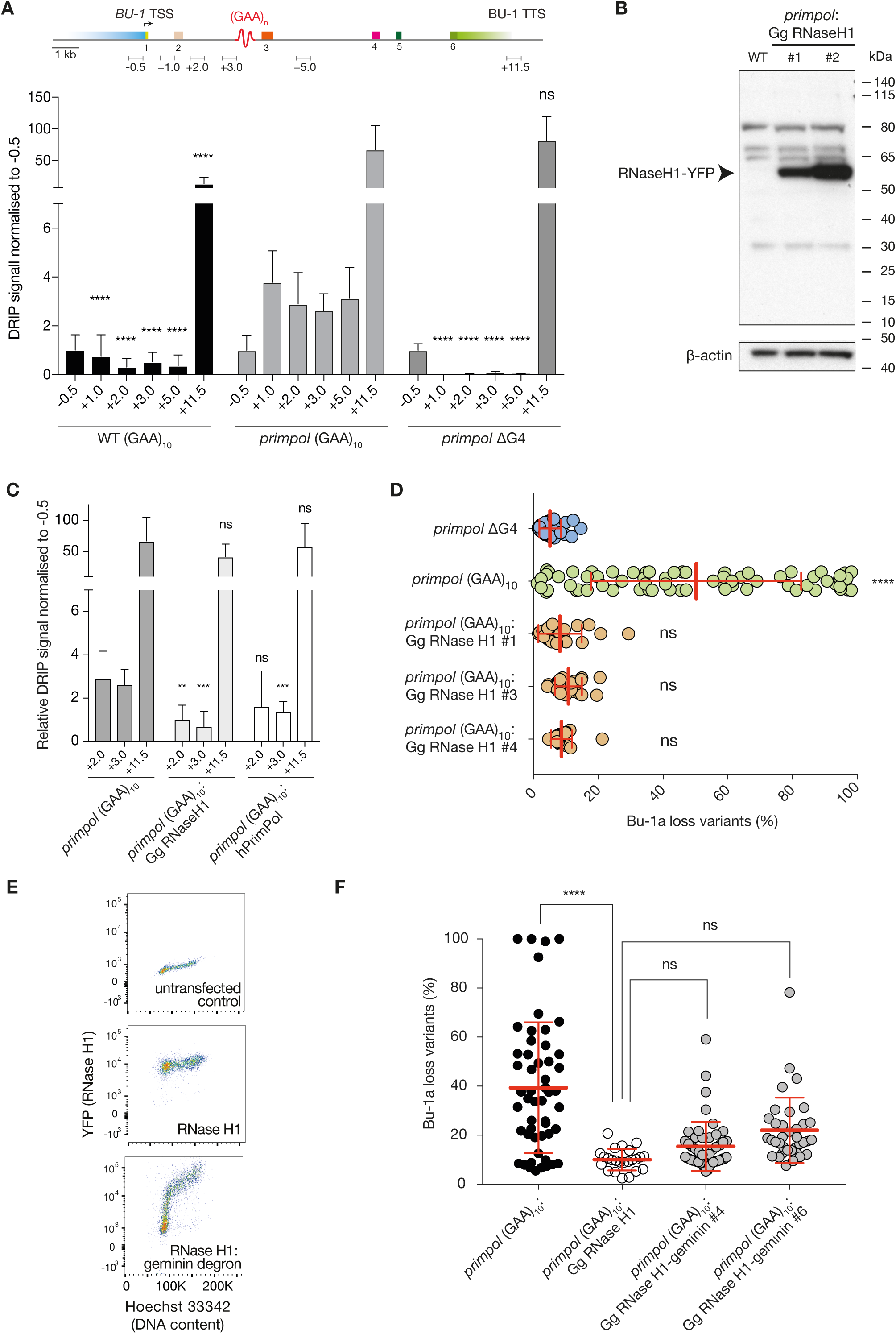
R-loops promote (GAA)_10_-dependent epigenetic instability of *BU-1*. A. DRIP-qPCR analysis reveals accumulation of R-loops across the *BU-1* locus in *primpol* cells. The DRIP signal was calculated as enrichment over RNaseH-treated samples and was normalised to −0.5 kb amplicon. The mean ± SD for three biological replicates is presented. An unpaired t-test was used to compare differences between matched amplicons in *primpol BU-1A^(GAA)10^* and the other cell lines indicated. B. Representative western blot showing stable expression of chicken RNaseH1-YFP fusion protein in *primpol* cells. C. DNA:RNA hybrids in *primpol BU-1A^(GAA)10^*:GgRNaseH1 and *primpol BU-1A^(GAA)10^*:hPrimPol. An unpaired t-test on three biological replicates was used to compare differences to *primpol BU-1A^(GAA)10^* for each matched amplicon. D. Overexpression of chicken RNaseH1 prevents (GAA)_10_ induced *BU-1A* epigenetic instability in *primpol* cells. Fluctuation analysis was performed on three *primpol BU-1A (GAA)10* clones. One-way ANOVA was used to calculate the significance of differences in *BU-1* instability between *primpol BU-1A ^ΔG4^* and other cell lines. E. Example of geminin-tagged chicken RNaseH1-YFP expression. Different phases of the cell cycle were determined by flow cytometry and staining of DNA content in live cells by Hoechst 33342. The RNaseH1-geminin fusion protein is depleted in G1. In contrast, RNaseH1 levels remain stable irrespective of the phase of the cell cycle. F. Bu-1a fluctuation analysis of two independently derived *primpol BU-1A^(GAA)10^*: Gg RNaseH1-geminin degron clones. Statistical differences calculated using one-way ANOVA. For all panels, mean ± SD reported. **** p ≤ 0.0001, *** p ≤ 0.001, ** p ≤ 0.01, ns = not significant.

### R-loop formation is required for (GAA)_10_ to induce expression instability of BU-1 in primpol-deficient cells

We next asked whether formation of a replication block at (GAA)_10_ requires an R-loop. To address this, we overexpressed chicken RNaseH1, an enzyme that degrades R-loops (Stein & Hausen, 1969) and that we have previously shown can be stably expressed and active in DT40 cells (Romanello et al., 2016). YFP-tagged chicken RNaseH1 carrying a disrupted mitochondrial localisation sequence was stably expressed in *primpol BU-1A^(GAA)10^* cells (Fig 3b). Expression of this construct reduced the R-loop signal in the vicinity of the repeat (+3 kb) to a level comparable with the background signal (Fig 3c).

We next performed fluctuation analysis for the formation of Bu-1a loss variants in the RNaseH1-overexpressing *primpol BU-1A^(GAA)10^* cells. Strikingly, RNaseH1 completely prevented the formation of Bu-1a loss variants in three separate clones, an effect comparable to removing the (GAA)_10_ repeat itself (Fig 3d). This suggests that DNA:RNA hybrid formation makes a crucial contribution to the ability of (GAA)_10_ to induce *BU-1* expression instability.

Since the activity of PrimPol is intimately linked with replication, we hypothesised that specifically removing the R-loops in S phase would suppress (GAA)_10_-induced *BU-1* expression instability. We therefore expressed chicken RNaseH1 fused to a degron sequence from geminin, which ensures protein expression is restricted to S phase (Sakaue-Sawano et al., 2008) (Fig 3e). Expression of this construct was able to prevent instability of *BU-1* expression in *primpol BU-1A^(GAA)10^* (Fig 3f), confirming that R-loops present during S phase are indeed responsible for the (GAA)_10_-dependent destabilisation of *BU-1*. Together with the DRIP experiments, this result suggests that R-loop formation is not only a consequence of the (GAA)_10_ repeat but is necessary for it to become a replication impediment.

### R-loop stabilisation converts the (GAA)_10_ sequence into a replication impediment

The R-loop dependence of *BU-1* expression instability in *primpol* mutants led us to the prediction that the enforced stabilisation of R-loops might lead the (GAA)_10_ repeat to induce *BU-1* expression instability even in wild type cells. To achieve this, we overexpressed the 52 amino acid DNA:RNA hybrid binding domain (HBD) of human RNaseH1, a fragment previously shown by the Aguilera lab to co-localise with and stabilise DNA:RNA hybrids *in vivo* (Bhatia et al., 2014). We cloned the HBD in frame with mCherry separated by a flexible GSGSG linker. The resulting fusion protein could be stably expressed in DT40 cells as monitored by mCherry fluorescence and by Western blotting (Fig 4a & b). Expression of the HBD in cells lacking a structure-forming sequence at the +3.5 kb position of *BU-1A* (DT40 *BU-1A*^∆G4^) did not induce statistically significant destabilisation of *BU-1* expression compared with the control (Fig 4c). However, when the (GAA)_10_ repeat was present at the +3.5kb position we observed significantly greater expression instability. This observation provides further evidence that R-loops are causal in promoting a (GAA)_10_ motif to become a replication block.

**Figure 4.**
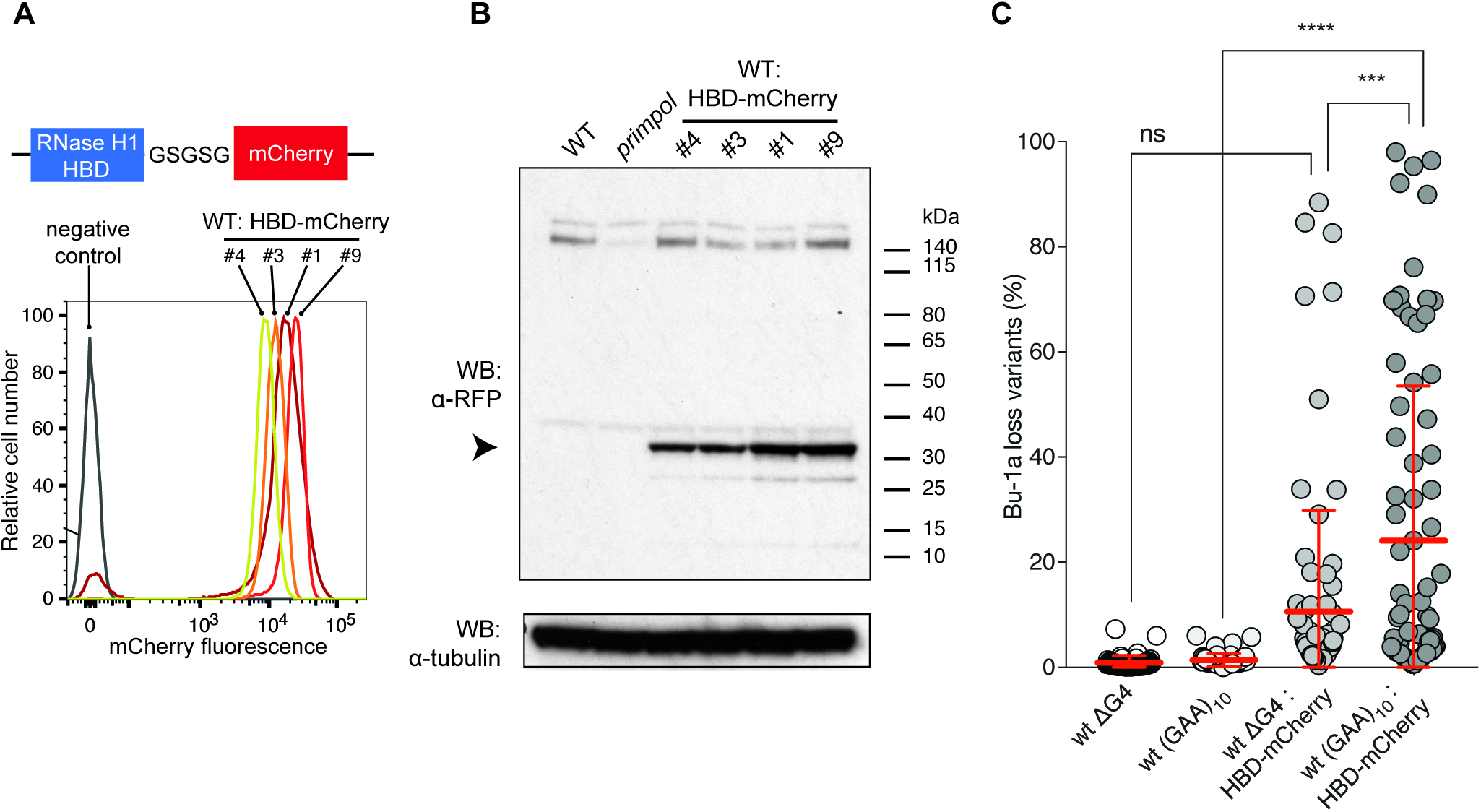
R-loop stabilisation induces epigenetic instability of *BU-1*. A. A diagram of the RNaseH1 hybrid binding domain (HBD) – mCherry fusion and flow cytometry expression profiles of the construct in four clones. B. Western blots showing stable overexpression of HBD-mCherry in the same four wild type DT40 clones as in A. C. Bu-1a fluctuation analysis of wild type cells expressing HBD-mCherry. The scatter plots pool results from at least two different clones with matched HBD-expression. Mean ± SD reported. **** p ≤ 0.0001,*** p ≤ 0.001, ns = not significant; one-way ANOVA.

### PrimPol curtails R-loop formation during S phase

As the activity of PrimPol is linked with replication, we next asked if the accumulation of R-loops at the *BU-1* locus of *primpol* cells (Fig 3a) occurs during S phase. Wild type *BU-1A^(GAA)10^* and *primpol BU-1A^(GAA)10^* cells were synchronised in G1 by double thymidine block, with cells at different phases of S phase sampled upon release (Fig EV6a) and steady-state R-loop dynamics monitored using DRIP (Fig 5a). In wild type cells, the highest levels of DNA:RNA hybrids occurred in G1 and G2 phases, most likely associated with transcription, while they remained relatively sparce during S phase. In contrast, R-loops in *BU-1* of *primpol* cells were significantly elevated at the early stages of S phase relative to wild type. Using the published replication timing map for *BU-1* (Schiavone et al., 2014) we estimate that the locus is replicated within 2 hours of entry into S phase, which corresponds to the time points at which we observe an increase in R-loop signal.

**Figure 5.**
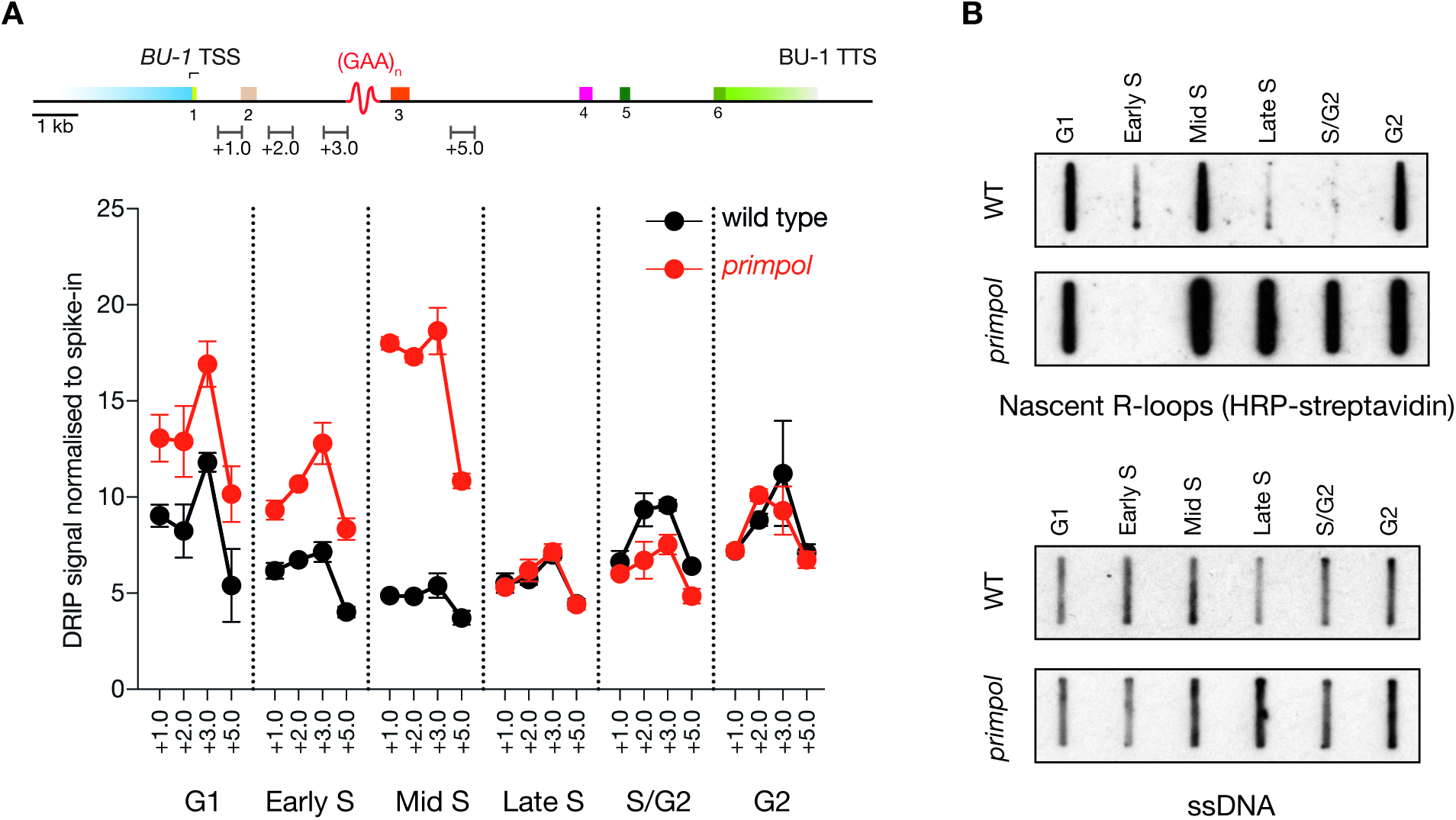
PrimPol suppresses R-loop formation in S phase. A. Cell cycle DRIP-qPCR for R-loops around the engineered +3.5 (GAA)_10_ repeat in *BU-1.* The location of the qPCR amplicons is indicated in the map at the top of the Fig. Cells in each phase of the cell cycle were spiked in with human TK6 cells in the same ratio to act as the calibrator. The *BU-1* DRIP signal was normalised to R-loop positive sites in human cells (*RPL13A*, *TFPT* and *FBXL17*). See Fig EV6a for cell cycle synchronisation profiles. Black: wild type; red: *primpol*. Error bars = SD. B. Nascent R-loop assay (See Fig EV6b for experimental scheme). Newly formed DNA:RNA hybrids were labelled with a 30’ pulse of 4-thiouridine at the indicated cell cycle stage (see also Fig EV6a). Upper panel: immunoblot of 2 μg of biotinylated nucleic acids detected with streptavidin-HRP; Lower panel: ssDNA loading control generated by DTT stripping of the blot.

To explore whether PrimPol suppresses R-loop formation in S phase, we used metabolic labelling of nascent RNA with 4-thiouridine (4-sU) to examine active DNA:RNA hybrid formation in different phases of the cell cycle. DNA containing R-loops was gently extracted, and free cytoplasmic and nuclear RNA was degraded with RNaseA in high salt leaving behind the RNA moiety of R-loops. Surviving 4-sU-labelled RNA was biotinylated and its relative abundance analysed by immunoblotting with streptavidin-HRP (Fig EV6b). Using this analysis, which is independent of the S9.6 antibody, we observed that, in wild type cells, the majority of R-loop formation is seen in the G1 and G2 periods of the cell cycle when co-transcriptional R-loop formation would be expected (Fig 5b). Interestingly, R-loop formation also increases transiently in early S phase, but then diminishes. In contrast, *primpol* cells exhibit a sustained DNA:RNA hybrid signal throughout S phase. Thus, the increased R-loops we observe in *primpol* cells result from unscheduled DNA:RNA hybrid formation during S phase. Extending from the experiments showing that R-loop accumulation in *BU-1* is potentiated by a repriming defect (Fig 3a), the unscheduled R-loop formation in S phase can be interpreted as the consequence of failure to restrict the length of single stranded DNA gaps produced by interruptions of DNA polymerisation as S phase progresses.

### PrimPol suppresses R-loop formation in the vicinity of secondary structure-forming sequences throughout the genome

To explore whether our observations at *BU-1* could be extended to the whole genome, we performed quantitative high-throughput sequencing of S9.6 immunoprecipitated DNA (DRIP-Seq) from wild type and *primpol* DT40 cells. To allow quantitation of the DRIP signal the DT40 cells were spiked with a fixed proportion of *Drosophila* S2 cells to provide an internal control (Orlando et al., 2014). Following read alignment and peak calling, peaks in the wild type and *primpol* samples were normalised to the mean number of *Drosophila* reads. This revealed a highly significant increase in the height of the DRIP peaks in *primpol* cells (Fig 6a). Between wild type and *primpol*, 84% of peaks were shared suggesting that the loss of PrimPol does not result in the appearance of new peaks, but that for any given peak there is a greater DRIP signal, suggesting a higher steady-state level of R-loops (Fig 6b). 41% of DRIP peaks overlapped with genes and 83% of genes with DRIP peaks were shared between wild type and *primpol*. We next asked if genes with DRIP peaks are enriched for H-DNA motifs. Using the ‘Triplex’ R package (Hon et al., 2013), sequences with H-DNA-forming potential were identified as overlapping just under 8% of all genes. The subset of genes harbouring DRIP peaks was significantly enriched for these sequences, with *c.* 15% of these genes overlapping sequences with H-DNA-forming potential (Fig 6c). Within this set of genes there was a significant increase in peak height in the *primpol* mutant (Fig 6d).

**Figure 6.**
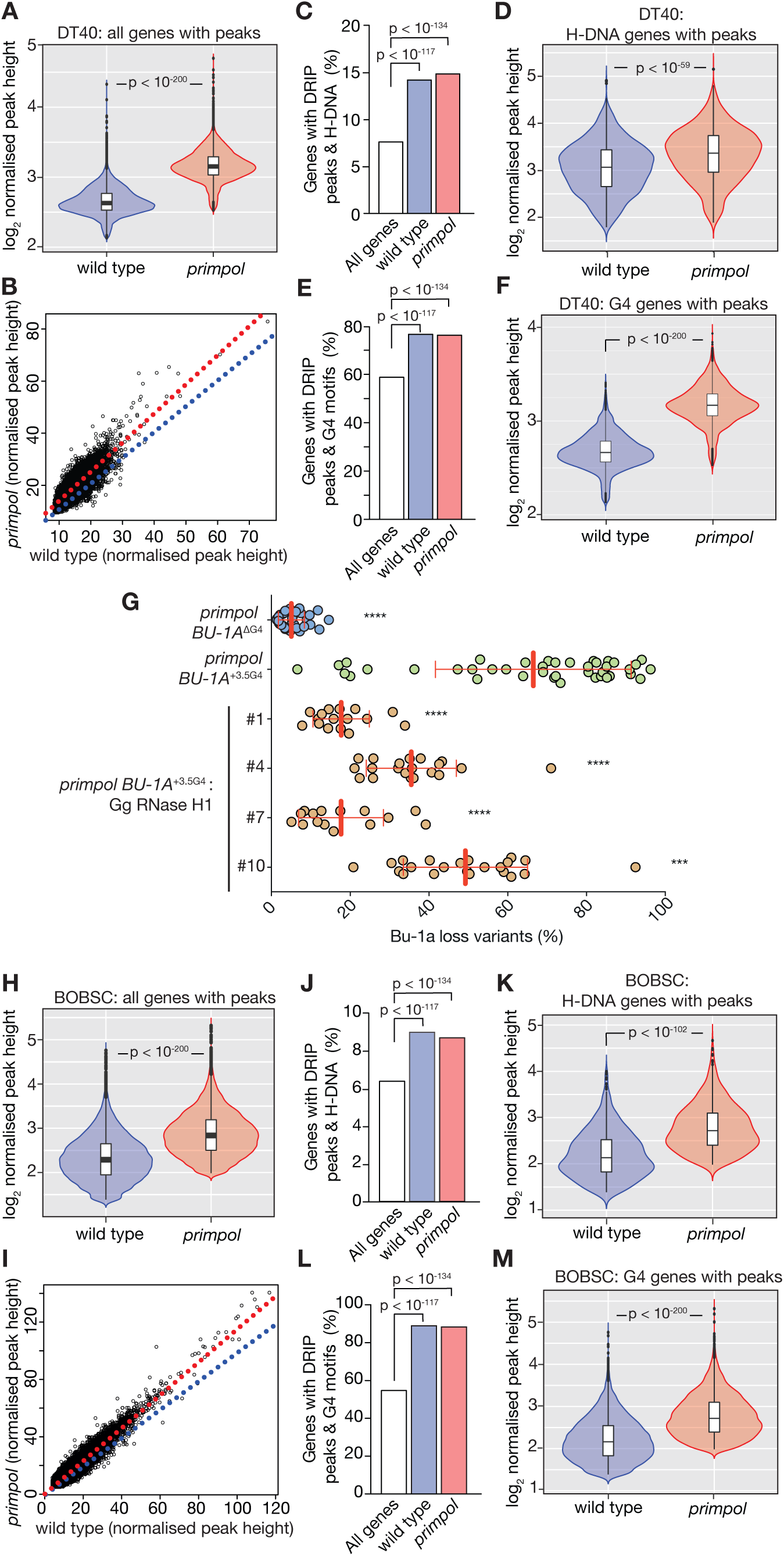
PrimPol suppresses R-loop formation in association with DNA secondary structure-forming sequences across the genome. A. DRIP peak heights in wild type and *primpol* DT40 normalised to *Drosophila* S2 spike-in. n (wild type) = 41445; n (*primpol*) = 48648. B. Correlation of normalised DRIP peak heights in the overlapping peaks between wild type and *primpol*. Blue line = 1:1 correlation; red line = linear regression through data. C, E. Correlation between H-DNA or G4 motifs ([G_3-5_N_1-7_]_4_) forming sequences and all genes (white bar) and genes with DRIP peaks and wild type (blue) and *primpol* cells (red) is shown in C and E, respectively. D, F. Normalised DRIP peak heights in the genes identified as associating with H-DNA and G4 motifs is shown in D and F, respectively. G. Overexpression of chicken RNaseH1 reduced G4-induced *BU-1* epigenetic instability in *primpol* cells. Fluctuation analysis was performed on four *primpol BU-1A^+3.5G4^* clones. One-way ANOVA was used to calculate differences in *BU-1A* instability between *primpol BU-1A^+3.5G4^* and other cell lines, including *BU-1A^∆G4^* (****p ≤ 0.0001, *** p ≤ 0.001). H. RNA-DIP peak heights in wild type and *primpol* BOBSC iPS cells normalised to DT40 spike-in. n (wild type) = 32740; n (*primpol*) = 33721. I. Correlation of normalised RNA-DIP peak heights in the overlapping peaks between wild type and *primpol*. Blue line = 1:1 correlation; red line = linear regression through data. J, L. Correlation between H-DNA or G4 motifs ([G_3-5_N_1-7_]_4_) and all genes (white bar) and genes with RNA-DIP peaks and wild type (blue) and *primpol* cells (red) is shown in J and L, respectively. K, M. Normalised RNA-DIP peak heights in the genes identified as associating with H-DNA and G4 motifs is shown in K and M. Data information. P values calculated with Mann-Whitney U test (A, D, F, H, K and M) and hypergeometric test (C,E,J and L). In violin plots, bar = median; box = interquartile range.

Our previous work has shown that G4s are able to induce similar epigenetic instability to the (GAA)_10_ repeat that has been the focus of this study. Further, G4 motifs have been linked to R-loop formation (Duquette et al., 2004). We therefore used the expression [G_3-5_N_1-7_G_3-5_N_1-_ 7G_3-5_N_1-7_G_3-5_] (Huppert & Balasubramanian, 2005) to identify G4 motifs in the chicken genome. G4 motifs were identified to overlap with 59% of all genes, while 76% of genes with DRIP peaks overlapped motifs, a significant enrichment (Fig 6e) and, again, the heights of the peaks in the *primpol* data set were significantly increased (Fig 6f).

The striking increase in steady state R-loop accumulation in genes containing G4 motifs and our previous demonstration that G4 motifs also potently destabilise *BU-1* expression in *primpol* cells (Schiavone et al., 2016) prompted us to ask if R-loops also contribute to promoting G4 motifs to become replication impediments. Forcing RNaseH1 expression resulted in a highly significant reduction in *BU-1* instability induced by the natural +3.5 G4 in four separate RNaseH1-expressing clones (Fig 6g), demonstrating that R-loop formation increases the probability of this G4 forming a significant replication impediment, but that it is not essential for it to do so.

Finally, we asked whether loss of PrimPol affected R-loop levels in human cells. PrimPol was disrupted using CRISPR/Cas9 editing in the induced pluripotent stem cell line BOBSC (Yusa et al., 2011). Genome wide R-loops were isolated with S9.6 immunoprecipitation followed by a modified sequencing protocol, RNA-DIP, that monitors the RNA moiety within the DNA:RNA hybrids. The analysis, normalised by spiking with DT40 cells, revealed a very similar picture to that seen in DT40. A highly significant increase in peak height in the *primpol* mutant (Fig 6h) reflected an increase in the RNA-DIP signal at existing peaks, with 66% of wild type and *primpol* peaks overlapping (Fig 6i). As in DT40, the genes containing peaks were associated with H-DNA and G4 motifs (Fig 6j & l) but in both cases the mean peak height was significantly higher in *primpol* cells (Fig 6k & m) demonstrating that loss of PrimPol also increases R-loop formation in the vicinity of DNA secondary structures in human cells.

## Discussion

### A requirement for PrimPol reveals that (GAA)_10_ forms a replication impediment

The (GAA)_10_ repeat upon which this study has focussed is typical of widespread short tandem repeats found throughout vertebrate genomes (Willems et al., 2014). Repeats of this length have not been previously linked to detectable disturbances in replication or transcription *in vitro* (Bidichandani et al., 1998, Ohshima et al., 1998) despite their potential to form triplex structures at physiological pH (Potaman et al., 2004). Our previous work supports a model in which instability of *BU-1* expression induced by G4s results from uncoupling of DNA unwinding from leading strand DNA synthesis (Sarkies et al., 2010, Schiavone et al., 2014, Šviković & Sale, 2017), and have shown that this uncoupling can be reduced by repriming (Schiavone et al., 2016). We now show that the repriming function of PrimPol is frequently deployed at short tandem repeats present throughout the genome as exemplified by the engineered (GAA)_10_ repeat. Consequently, loss of PrimPol causes widespread replication defects (Bianchi et al., 2013, Mouron et al., 2013).

### The nature of the replication impediment formed by (GAA)_10_

(GAA)_n_ repeats, in common with other polypurine-polypyrimidine tracts, are capable of forming triplex secondary structures in which a third strand anneals through Hoogsteen base pairing. This tendency has been linked to the detrimental effect of long GAA repeats on transcription elongation (Bidichandani et al., 1998, Punga & Buhler, 2010) through the trapping of transcribing RNA polymerase II (Grabczyk & Fishman, 1995). However, the very act of transcription also promotes formation of secondary structures (Kouzine et al., 2017, Lilley, 1980), including triplexes (Grabczyk & Fishman, 1995, Kouzine et al., 2004). This is likely driven by structure formation releasing the negative supercoiling generated in the wake of translocating RNA polymerase (Levens et al., 2016, Liu & Wang, 1987). Similar topology-induced structure formation could also contribute to leading strand secondary structure formation behind the replicative helicase (reviewed in Kurth et al., 2013, Yu & Droge, 2014).

We show that (GAA)_10_ requires an RNaseH1-sensitive R-loop in order to create a replication impediment that requires PrimPol-dependent repriming. It is well established that formation of DNA:RNA hybrids coincides with the sequences able to adopt non-B DNA (Duquette et al., 2004, Grabczyk et al., 2007) and is favoured in a negatively supercoiled DNA template (Roy et al., 2010). R-loops have been implicated as a major factor in the severity of head-on collisions between the replication and the transcriptional machinery (Hamperl et al., 2017). However, a direct head-on collision with transcribing RNA polymerase is likely to halt the entire replisome (Pomerantz & O’Donnell, 2010), precluding the displacement of parental nucleosomes caused by the functional uncoupling between the replicative helicase and DNA synthesis. It is difficult to reconcile this type of stall with the involvement of PrimPol. Specifically, the DNA and RPA binding activities of the enzyme suggests the transient formation of ssDNA, which most likely arises as a result of functional uncoupling of replicative helicase from the replicative polymerases, and which is the basis for *BU-1* expression instability.

How then can a (GAA)_n_ repeat generate the uncoupling of DNA unwinding and leading strand DNA synthesis necessary to induce transcriptional instability of *BU-1*? We propose that transcription of the (GAA)_10_ repeat generates an R-loop (Fig 7). During replication, the approaching replicative helicase traverses the transcription complex by displacing the RNA polymerase (Pomerantz & O’Donnell, 2010) or by reorganising the helicase itself (Huang et al., 2013, Vijayraghavan et al., 2016). Biophysical calculations show that DNA:RNA hybrids are sufficiently thermodynamically stable to survive the accumulation of positive supercoiling generated ahead of the replicative helicase (Belotserkovskii et al., 2013). Since the eukaryotic replicative helicase tracks on the leading strand (Douglas et al., 2018), we suggest that the DNA:RNA hybrid could remain intact on the lagging strand during passage of the helicase. Behind the replicative helicase, the persistent lagging strand DNA:RNA hybrid may retrap the purine-rich leading strand through triplex formation. The resulting R:R•Y triplex would then block leading strand synthesis (Samadashwily & Mirkin, 1994). This model is consistent with the observation that the depletion of DNA:RNA hybrids through overexpression of RNaseH1 completely abolishes (GAA)_10_-dependent *BU-1* expression instability in PrimPol-deficient cells. An alternative explanation for the creation of a leading strand impediment is the formation of a DNA triplex stabilised by an adjacent DNA:RNA hybrid, of the form proposed by Grabczyk and Fishman (1995). In either event, continued helicase activity would result in exposure of ssDNA ahead of the stalled replicative polymerase, which through being bound by RPA promotes the recruitment of PrimPol. Repriming then allows DNA synthesis to remain coupled to unwinding leaving the triplex in a small gap to be disassembled post-replicatively (Fig 7).

**Figure 7.**
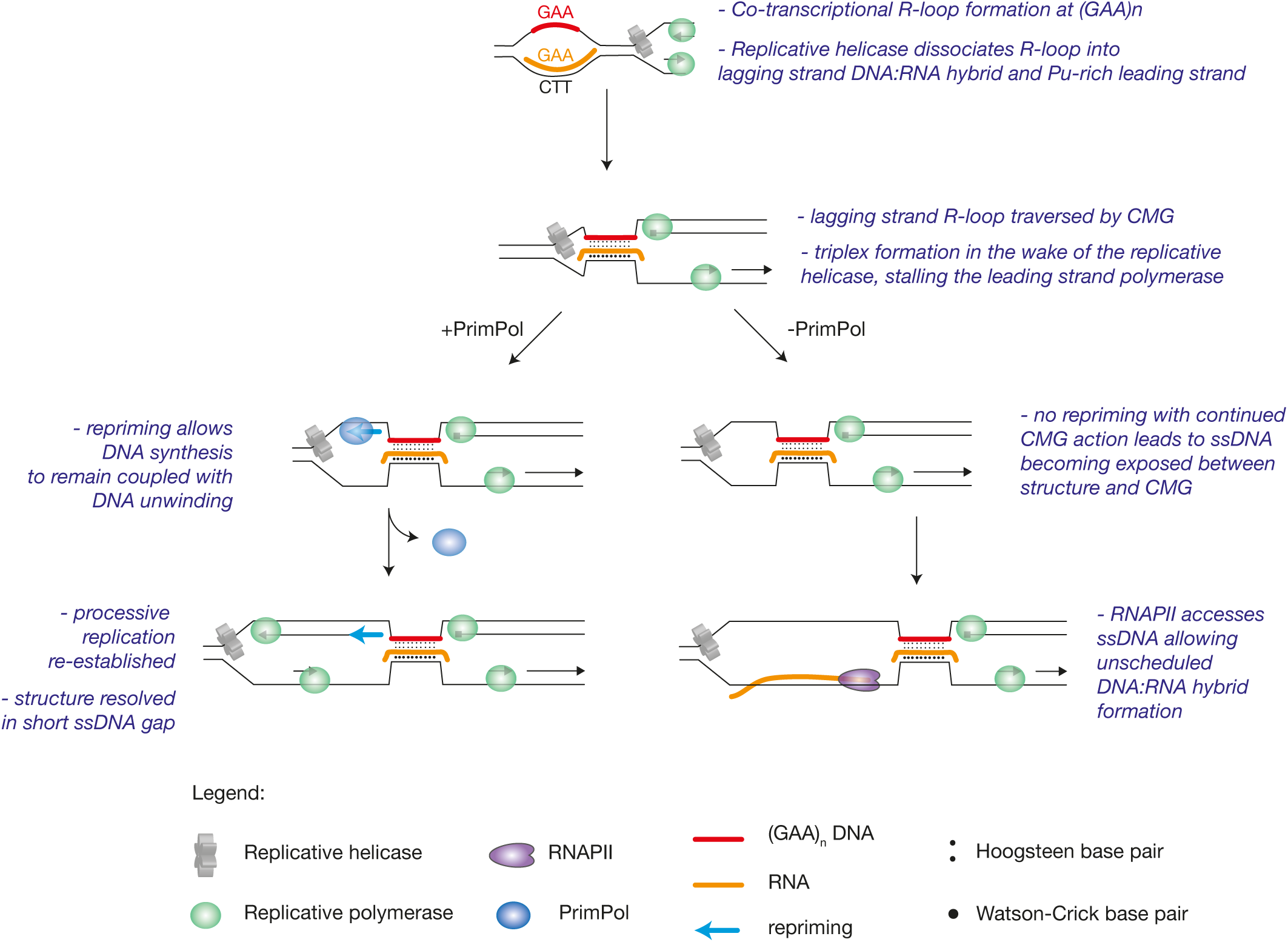
A model for how PrimPol suppresses S phase R-loops in the vicinity of DNA secondary structures. During transcription, an R-loop is formed at the GAA tract, which the replicative helicase dissociates during replication into purine-rich leading strand and DNA:RNA hybrid on the lagging strand. In the wake of the replicative helicase, triplex is formed between DNA:RNA hybrid and purine-rich sequence, blocking the leading strand polymerase. In wild type cells, PrimPol is recruited to the single stranded gap, where repriming allows DNA synthesis to remain coupled to fork progression. In the absence of Primpol, however, the activity of replicative helicase exposes long stretches of ssDNA. RNA polymerase II gains access to such template, leading to unscheduled DNA:RNA hybrid formation.

### Loss of PrimPol-mediated repriming at structured DNA promotes S phase R-loop accumulation

In the absence of PrimPol, continued unwinding of the parental duplex by the replicative helicase in the context of a continued stalling of DNA synthesis would create a more extensive region of ssDNA. That this results in a net increase in DNA:RNA hybrids in the vicinity of structure-forming sequences (Fig 3, 5b & 6) suggests that there is greater ssDNA exposure on the lagging, as well as the leading strand, which may explain the increase in nascent transcription during S phase (Fig 5a). These observations imply that RNAPII continues to transcribe despite its template remaining single stranded. This idea is consistent with both biochemical (Kadesch & Chamberlin, 1982) and *in vivo* reports (Michelini et al., 2017, Ohle et al., 2016) demonstrating that DNA:RNA hybrid formation can occur at ssDNA generated by resection of DNA ends at double-stranded DNA breaks. We suggest that ssDNA generated as a consequence of helicase-polymerase uncoupling, unmitigated by repriming, could also act as a substrate for unscheduled RNAPII transcription and DNA:RNA hybrid formation (Fig 7).

The results we present here establish two important mechanistic points concerning the relationship between R-loops and impeded replication. First, we show that R-loops are able to promote short sequences with structure-forming potential to become replication impediments, requiring the repriming activity of PrimPol to maintain their processive replication. Second, failure to reprime at these sequences increases R-loop formation. We suggest this is due to failure to prevent the exposure of excessive single stranded DNA during S phase. This relationship between structure-forming DNA and repriming is seen at a significant fraction of R-loops detected across the genome, suggesting that repriming plays an important role in allowing cells to manage the complex challenges created by the interaction of transcription and replication.

## Materials & Methods

### Cell culture and transfection

DT40 cell culture, cell survival assays, fluctuation analysis for generation of Bu-1 loss variants (Fig EV1), and genetic manipulation of the *BU-1* locus were performed as previously described (Schiavone et al., 2014, Simpson & Sale, 2003, Simpson & Sale, 2006). Drosophila S2 cells were grown in Insect-XPRESS Protein-Free Insect Cell medium with L-glutamine (Lonza), supplemented with 1% penicillin-streptomycin at 27 °C, ambient CO_2_ with 105 rpm agitation. BOBSC human induced pluripotent stem (hiPS) cells (Yusa et al., 2011) were cultured feeder-free on dishes coated with Vitronectin XF (07180, Stem Cell Technologies) in Essential 8 Flex media (A2858501, ThermoFisher Scientific) at 37 °C and 5% CO_2_. Cells were split 1:10 −1:15 every 3 – 4 days depending on confluence. All cells tested negative for mycoplasma.

### CRISPR-Cas9 mediated gene disruption in human cells

Guide RNA sequences used for disrupting *PRIMPOL* in BOBSC iPS cell lines are listed in the Appendix. Each gRNA sequence was cloned into pX458 (Ran et al., 2013). A targeting construct carrying puromycin selection marker was constructed by Gibson assembly using PCR amplified 5’ and 3’ homology arms (primer sequences in the Appendix). Equimolar amounts of targeting construct, gRNA expression vectors and Cas9 expression vectors were delivered by Amaxa electroporation. Puromycin resistant clones were genotyped by PCR and Sanger sequencing (Shen et al., 2014).

### Molecular cloning and transgene constructs

For transgene expression, cDNA was cloned in frame with fluorescent protein (mCherry or YFP) in the polylinker of pXPSN2 (Ross et al., 2005). The expression module was released with SpeI digestion and subcloned into pBluescript based vectors containing a loxP flanked puromycin or blasticidin S selection cassettes (Arakawa et al., 2001), which were transfected into DT40 via electroporation. Primers used for molecular cloning are listed in the Appendix. The hPrimPol-YFP, hPrimPol [AxA]-YFP and hPrimPol [ZfKO]-YFP constructs for complementation of *primpol* DT40 were previously described (Schiavone et al., 2016). cDNAs for PrimPol RPA binding mutants (RBM-A and RBM-B) were PCR amplified from previously described vectors (Guilliam et al., 2017) with primers listed in the Appendix. Similarly, chicken RNaseH1 (lacking the mitochondrial localisation sequence) was PCR amplified from DT40 cDNA and cloned in frame with YFP on the C-terminus. For cell cycle-regulatable RNaseH1, human Cdt1 [30-120] and geminin [1-110] fragments were PCR amplified from pLL3.7m-Clover-Geminin(1-110)-IRES-mKO2-Cdt(30-120) from the Fucci4 system (Bajar et al., 2016) and cloned in frame to the C-terminus of chicken RNaseH1ΔMLS-YFP. The hybrid binding domain was amplified from human cDNA using primers previously published (Bhatia et al., 2014) and fused to C-terminal mCherry via flexible GSGSG linker. Uninterrupted GAA tracts were created using a previously developed strategy (Scior et al., 2011), see Fig EV2 for details. The acceptor plasmid was created by inserting a linker containing a (GAA)_20_ repetitive tract flanked by recognition sites for the restriction enzymes BbsI, BsmBI and NcoI into pBluescript SK(+). Plasmids containing the repeats were transformed into a strain of *E. coli* lacking the SbcCD nuclease, DL733 (Connelly et al., 1997), a kind gift of David Leach, to avoid excision of secondary structures caused by the repeat. The confirmed uninterrupted repeats with minimal flanking sequence were released by MluI digest, subcloned into the BU-1 targeting construct and screened for orientation by Sanger sequencing.

### Chromatin immunoprecipitation (ChIP)

Chromatin immunoprecipitation was performed as previously described (Schiavone et al., 2014). Solubilised chromatin was diluted and immunoprecipitated overnight using the following antibodies: histone H3 (ab1791, Abcam), H3K4me3 (9727, Cell Signaling), and Normal rabbit IgG (2729, Cell Signaling). Primer sequences are provided in the Appendix. Enrichment was normalized to H3 and reported relative to wild-type.

### Bisulphite sequencing

bisulphite conversion of genomic DNA was performed using EZ DNA Methylation-Gold Kit (Zymo Research) as per the manufacturer’s instructions. PCR was performed with ZymoTaq (Zymo Research) and primers compatible with bisulphite-converted DNA (Appendix) for 40 cycles according to manufacturer’s instructions. PCR products were purified, digested with NotI and SacI, size selected by gel extraction and cloned into pBK CMV. Individual bacterial colonies were sequenced by GATC.

### DNA:RNA immunoprecipitation (DRIP)

After fractionation and lysis of nuclei (see Slot-blot detection of genomic DNA:RNA hybrids), genomic DNA containing DNA:RNA hybrids were digested overnight with a restriction enzyme cocktail containing BamHI, NcoI, PvuII, ApaLI, and NheI, yielding and average fragment size of 1 kb. Samples were subsequently diluted to 5 ml with IP dilution buffer (16.7 mM Tris HCl pH 8.0, 1.2 mM EDTA, 167 mM NaCl, 1.1% Triton X-100, 0.01% SDS), pre-cleared for two hours with 30 μl Protein-G Sepharose beads (Dharmacon) and immunoprecipitated with 10 μg S9.6 antibody or with mouse IgG isotype control (10400C; Invitrogen) overnight at 4 °C. Subsequent steps are essentially the same as for ChIP. One third of the digested material was subsequently treated with 25 U of RNase H (NEB, M0297) overnight at 37 °C. The signal across *BU-1* locus was normalized to RNaseH background signal and baselined to 28S rDNA.

### Cell cycle synchronization and 2D cell cycle analysis

G1 phase synchronization of DT40 cells was achieved by double thymidine block. Cells were treated overnight with 2 mM thymidine, released for 9 hours, and again treated with thymidine overnight, after which cells were released into medium containing 0.2 μM nocodazole. Upon release, cells in different cell cycle phases were harvested to be analysed or pulse labeled with BrdU. 5 - 10×10^6^ DT40 cells were pulse labelled with 50 μM BrdU for 30 minutes in complete medium at 37 °C. BrdU staining was performed as previously described (Frey et al., 2014).

### 4-thiouridine (4sU) metabolic labelling of nascent DNA:RNA hybrids

4-thiouridine was used to label nascent RNA during different stages of cell cycle. 75-100 million DT40 cells were resuspended in 10 ml of complete medium supplemented with 100 μM 4sU and incubated for 30 minutes at 37 °C. 4sU incorporation was terminated by adding ice cold PBS, after which cells were lysed in TE buffer supplemented with 1% SDS and 0.2 mg/ml Proteinase K. Nucleic acids were purified with phenol:chloroform:isoamyl alcohol and digested with KpnI, EcoRI and BamHI overnight at 37°C, followed by treatment with 100 μg RNaseA in 0.5 M NaCl. RNaseA in high-salt inactivates its RNaseH-activity (Halasz et al., 2017). DNA was purified again with phenol:chloroform and resuspended in 10 mM Tris-HCl pH 8.0. 4sU-labelled RNA moiety of DNA:RNA hybrids were biotinylated with thiol-specific reaction using EZ-Link HPDP-Biotin (Thermo Scientific) as described previously (Fuchs et al., 2015). Free biotin was removed by chloroform:isoamyl alcohol (24:1 v/v). Nucleic acids were bound to Hybond-XL membrane using slot blot apparatus, crosslinked and blotted with HRP-Conjugated Streptavidin (N100, Thermo Scientific). As DMF is a known DNA denaturation agent (Cortadas & Subirana, 1977, Herskovits et al., 1961), generated ssDNA was used as an internal loading control. Biotin was stripped for one hour in 100 mM dithiothreitol at room temperature, and the membrane incubated with anti-ssDNA antibody.

### Chromatin associated RNA (ChrRNA) extraction

RNA associated with the chromatin was extracted as described previously (Nojima et al., 2016). Briefly, DT40 cells were lysed in HLB+N (10mM Tris-HCl (pH 7.5), 10 mM NaCl, 2.5 mM MgCl_2_ and 0.5% (vol/vol) NP-40) and passed through a 10% sucrose cushion. Nuclei were then resuspended in 125 μl NUN1 buffer (20 mM Tris-HCl (pH 7.9), 75 mM NaCl, 0.5 mM EDTA and 50% (vol/vol) glycerol), mixed with 1.2 ml NUN2 (20 mM HEPES-KOH (pH 7.6), 300 mM NaCl, 0.2 mM EDTA, 7.5 mM MgCl_2_, 1% (vol/vol) NP-40 and 1 M urea) vortexed vigorously and spun for 10 minutes at 16,000 g. Chromatin pellets were digested twice with DNase I (NEB, M0303) and Proteinase K. ChrRNA was extracted with QIAzol (QIAGEN) and converted to cDNA using QuantiTect Reverse Transcription Kit (QIAGEN) as recommended by the manufacturer. The enrichment of RNA across the *BU-1* locus was analysed by qPCR, and the signal normalized to GAPDH.

### DRIP-Seq

Sample preparation for DRIP-Seq was essentially performed as described above, but with some minor changes. All the samples were spiked (Orlando et al., 2014) with the same batch of Drosophila S2 cells in 1:4.2 ratio to DT40 cells. Digested DNA was treated with 100 μg RNase A in the presence of 0.5 M NaCl for 2 h at 37°C. Elution from magnetic beads was performed for 1 h at 37 °C in 300 μl elution buffer supplemented with 0.1 mg/ml RNase A. To ensure complete elution, 10 μg Proteinase K was added and incubated further 90 minutes at 37 °C. DNA was purified by phenol:chloroform:isoamyl alcohol extraction, quantified with Qubit dsDNA HS Assay Kit (Invitrogen), diluted with ultra-pure water to 55 μl and sheared with Covaris M220 Focused-ultrasonicator and Holder XTU to average size of 300 bp in microTUBE-50 AFA. DNA libraries were built using NEBNext Ultra II DNA Library Prep Kit (New England Biolabs) as per the manufacturer’s instructions.

### RNA DIP-Seq

Between 60-100 million human cells were spiked in with DT40 cells (1/10 ratio), harvested, washed with cold PBS and nuclei isolated by lysing cells in HLB+N and passing it through a 10% sucrose cushion. Collected nuclei were lysed overnight in with NLB (25 mM Tris-HCl (pH 7.5), 1% SDS, 5 mM EDTA, 0.125 mg/ml Proteinase K) with agitation at 37°C. Nucleic acids were purified with 1 M potassium acetate (pH 5.5), precipitated and treated with of RNaseI (Ambion, AM2294) per 90 μg of DNA (15 minutes at 37°C) to degrade soluble RNA molecules. DNA was purified with phenol:chloroform, diluted with the IP dilution buffer and sheared with a Bioruptor plus (Diagenode) to average size of 300 bp. DNA:RNA hybrids were immunoprecipitated with S9.6 mAb (1 μg antibody for each 2 μg of DNA) overnight. Immunocomplexes were captured with Protein G beads and washed as for ChIP and DRIP preparation. DNA:RNA hybrids were eluted by incubating the sample with Proteinase K for 2 hours at 42°C. Nucleic acids were cleaned up with phenol:chloroform:isoamylalchohol, precipitated with glycogen and resuspended in water, denatured for 5 minutes at 90°C and immediately placed on ice. DNA moiety of DNA:RNA hybrids were removed with 4U DNase I for 30 minutes at 37°C. RNA was extracted with QIAzol, precipitated overnight and dissolved in RNase-free water. Strand-specific Illumina-compatible libraries were prepared with NEBNext Ultra II Directional RNA Library Prep Kit (NEB, E7760) with 100 ng input. Libraries were quality checked as before and sequenced on a NextSeq500 (Illumina).

### Quantification and statistical analysis of deep sequencing data

DRIPseq libraries were sequenced on an Illumina HiSeq 4000, RNA-DIP libraries were sequenced on an Ilumina NextSeq. Reads were trimmed and quality filtered using trim galore (version 0.4.4) (https://www.bioinformatics.babraham.ac.uk/projects/trim_galore/), then aligned to genomes with bowtie2 (version 2.26) (Langmead & Salzberg, 2012) using default settings. DT40 reads were aligned to Ggal 5.0 and Dmel r6.18, BOBSC reads were aligned to GRCh38 and Ggal 5.0. Alignments were filtered for uniquely matching reads and separated into sample and spike-in. Peaks were called on filtered alignments using MACS2 (version 2.1.1.20160309) with the default settings and -g 1.87e9 (or -g hs for human) --broad (Feng et al., 2012). Peak heights were normalised to the read number of the spike-ins and compared using the Mann Whitney U test. Overlaps between peaks were calculated using bedtools2 closest (version 2.27.1) with default settings (Quinlan & Hall, 2010). Peaks were considered to be overlapping if at least 1bp overlapped. Sequences with H-DNA forming potential were identified with the Triplex R package (Hon et al., 2013). G4 motifs were identified using the Quadparser algorithm (Huppert & Balasubramanian, 2005) with the regex [G_3-5_N_1-7_G_3-5_N_1-_ 7G_3-5_N_1-7_G_3-5_]. Enrichment testing for secondary structures was performed using the hypergeometric test.

## Data Availability

Deep sequencing data has been deposited in the GEO repository (https://www.ncbi.nlm.nih.gov/geo/) with accession number GSE112747.

## Acknowledgements

We would like to thank Maria Daly and her team in the LMB flow cytometry facility for cell sorting, Toby Darling and Jake Grimmett of LMB Scientific Computing for assistance, Prof. David Leach (University of Edinburgh) for the gift of the SbcCD-deficient *E. coli* strain DL733, the CRUK Cambridge Institute Genomics Core for Illumina sequencing, Ludovic Vallier and the Wellcome Trust Sanger Institute for the BOBSC iPS line and Cara Eldridge for guidance on iPS line culture. The generation of the *primpol* BOBSC line was carried out under the auspices of the COMSIG consortium, supported by a Wellcome Trust strategic award (101126/B/13/Z). Finally, we thank all members of the Sale lab, Joe Yeeles, Martin Taylor and Pierre Murat for helpful discussions. S.Š. was funded by the LMB Cambridge International Scholarship and A.C. by a UKRI Innovation Fellowship. Work in the J.E.S. lab is supported by a core grant from the MRC to LMB (U105178808). The A.J.D. laboratory is supported by grants from the BBSRC: BB/H019723/1 and BB/M008800/1. T.A.G. was supported by a University of Sussex PhD studentship. The N.J.P. lab is supported by a Welcome Trust Investigator Award (107928/Z/15/Z) and an ERC Advanced Grant (339170).

## Author contributions

S.Š. and J.E.S. conceived the study and wrote the paper with input from all authors. S.Š. performed all the experiments. S.M.T.-W. and N.J.P. developed and with S.Š. performed the RNA DIP-seq experiments. T.A.G. and A.J.D. identified and created the PrimPol mutant cDNAs. A.C. and G.G. analysed the deep sequencing data.

